# The neural signature of magnitude integration between time and numerosity

**DOI:** 10.1101/2022.08.29.505731

**Authors:** I. Togoli, M. Fornaciai, E. Visibelli, M. Piazza, D. Bueti

**Author notes:** Corresponding author: Michele Fornaciai.

## Abstract

Magnitude dimensions such as time and numerosity are fundamental components of our visual experience, allowing us to understand the environment and interact with it. Different magnitudes are however not processed independently from each other, but show a relationship whereby the perception of one dimension depends on the others (“magnitude integration”). In this study, we use electroencephalography (EEG) to address whether such integration may arise from a shared brain processing stage where different dimensions are integrated together, or from independent parallel processes interfering with each other. In the experiment, participants judged either the average numerosity or duration of dynamic dot-array stimuli concurrently modulated in both dimensions. First, the behavioural results show a magnitude integration effect in both tasks, with duration affecting the judgement of numerosity and vice versa. The EEG results further show that both numerosity and duration significantly modulate event-related potentials at several distinct latencies. Crucially, however, we identified a significant interaction between numerosity and duration emerging in a specific latency window (360-460 ms) irrespective of the task performed by participants. In this latency window, the modulation of ERPs provided by the interfering magnitude dimension can be predicted by the strength of the behavioural bias. Our results thus support the idea of different magnitude dimensions converging onto a shared perceptual processing stage mediating their integration. Overall, our results demonstrate a clear electrophysiological signature of magnitude integration between numerosity and time, and provide new evidence for a shared representational system encompassing different magnitude dimensions.

## INTRODUCTION

Temporal and numerical information is ubiquitous in the environment that we face in our everyday lives. The brain processing of magnitude information thus plays an essential role in our understanding of the external world, and in guiding our interaction with the environment. Indeed, the different magnitudes of the environment allow the brain to structure a representation of the basic properties of a visual scene, like the number of objects or individuals, and the timing and duration of the events. Thus, perceiving magnitudes represents a fundamental process to navigate the environment and plan our actions according to the external stimuli.

Although time perception and numerosity perception are most often investigated in separate research lines, they are nevertheless characterized by an intrinsic link (e.g., Walsh, 2003). Indeed, different magnitude dimensions have been shown to interact with each other so that the processing of one dimension is influenced by another dimension. For example, an event involving a bigger stimulus or a more numerous set of objects is perceived as lasting longer in time compared to smaller stimuli or less numerous sets (e.g., Rammsayer & Verner, 2014; Togoli et al., 2021; Xuan et al., 2007). Vice versa, a longer duration can lead to an increase in the perceived size of a stimulus (Cai & Connell, 2015) or an increase in perceived numerosity (Javadi & Aichelburg, 2012; Togoli et al., 2021).

These interactions across different magnitude dimensions have led to the proposal of a generalized magnitude system (Walsh, 2003), encoding information from different dimensions with a similar neural code and integrating it to subserve behaviour. This process of *magnitude integration*, potentially increasing the consistency of perceptual representations by exploiting redundancies across dimensions (i.e., different magnitudes may covary in the external environment), would thus lead to the mutual biases observed in the experimental context. Regarding the potential neural correlates of this process, such a generalized magnitude system has been proposed to reside in the parietal cortex (Bueti & Walsh, 2009; Walsh, 2003), where a common network of brain areas would be responsible for the processing and integration of different magnitudes. Functional imaging results indeed provided evidence for a shared magnitude network located in parietal areas (Skagerlund et al., 2016), although a common neural code representing different dimensions has not been observed so far (Borghesani et al., 2019). Such a common network of brain areas however does not univocally support the existence of a truly generalized magnitude system; indeed, the processing and integration of different magnitude dimensions could be similarly carried out by overlapping but not converging magnitude channels (e.g., see for instance Harvey et al., 2013, 2015 for partially overlapping but different topographic maps of numerosity and object size).

When it comes to the nature of the magnitude integration effect itself (i.e., the bias provided by one magnitude to another), different interpretations have been proposed concerning perception or higher-level cognitive functions. Namely, it has been debated whether magnitude integration and its relative bias arises from a perceptual processing of magnitude information – thus effectively entailing a change in the phenomenological appearance of a stimulus dimensions as a function of the interfering magnitude – or whether the interaction occurs at a later cognitive stage. Indeed, one possibility is that the “magnitude integration” bias could not involve an actual integration occurring during perceptual processing, but rather a mnemonic interference between different magnitude representations stored in working memory (Cai et al., 2018; Cai & Wang, 2014). In this scenario, different magnitude dimensions would thus be processed in an independent fashion by distinct mechanisms, and then interfere with each other once stored in working memory. Moreover, different magnitude dimensions, as they usually share a similar “more than”/”less than” mapping, could even interfere at the time of response selection, explaining the magnitude integration effect in terms of a response bias (Yates et al., 2012). In a recent study from our group (Togoli et al., 2022), however, we demonstrated that a congruent, attractive magnitude integration effect selectively emerges only when two magnitudes are conveyed by the exact same object, ruling out the possibility of a response bias. Moreover, previous EEG results concerning the effect of symbolic numerical magnitude on duration processing showed an early effect more consistent with a perceptual rather than a decisional or mnemonic effect (Xuan et al., 2009). Nevertheless, whether the mutual magnitude integration biases arise from information converging onto a shared representational system, or from independent processes interacting only at a post-perceptual stage, remains unclear.

In the present study, we use electroencephalography (EEG) to investigate the neural signature of magnitude integration between time and numerosity, and address the existence of a shared processing stage mediating the interaction of these two dimensions. To do so, we used a numerosity and a duration classification task of dynamic dot-array stimuli modulated in duration and in average numerosity (i.e., the average number of dots presented across several briefly flashed arrays). Dynamic stimuli were specifically used to facilitate a mutual interaction between the two dimensions (Togoli et al., 2021), since static stimuli often yield a strong bias induced by numerosity on duration while not showing an effect in the opposite direction (e.g., Dormal et al., 2006). EEG data was then used to assess how changes in stimulus numerosity, duration, or their combination modulate the event-related potentials (ERPs) evoked by the stimuli in the two task conditions. In our hypothesis, the existence of a shared processing stage mediating the integration of different dimensions predicts that ERPs should be modulated by the interaction between numerosity and duration, with a temporal dynamic that should not depend on the task performed by participants. Alternatively, the bias could arise from processes unfolding in parallel and independently. In this case, the two processes might for instance interfere with each other without converging onto a shared processing stage, or interfere at a post-perceptual processing stage beyond the initial encoding of magnitude information. The existence of independent magnitude “channels” interacting via post-perceptual processes would thus predict the emergence of independent signatures of numerosity and duration processing, and modality-specific interaction effects (i.e., depending on which dimension is task-relevant and which is considered as interfering).

To preview, our behavioural results first show systematic magnitude integration biases from duration to numerosity and from numerosity to duration, albeit stronger in the former case. In terms of EEG results, we first show that the modulation of ERPs provided by numerosity and duration seems to not depend on the task performed by participants, suggesting that the neural signature of these magnitudes is not affected by attention and task-related factors. Moreover, the analysis of ERPs as a function of the different combinations of the two magnitudes show a consistent modulation of evoked activity induced by duration and/or average numerosity, across a series of both similar and distinct latency windows. Crucially, our results also show a specific latency window where ERPs are significantly modulated by the interaction between numerosity and duration, with a similar timing (360-460 ms after stimulus onset) irrespective of the task performed by participants. Moreover, further analyses across a series of latency windows show that, in each task condition, ERPs at 360-460 ms are significantly modulated by the interfering dimension, and that this modulation can be predicted from the strength of the behavioural bias.

## METHODS

### Participants

A total of 22 healthy adults with normal or corrected-to-normal vision and with a mean age of 23 years (SD = 2.82; age range = 18-31; 1 male) participated in the study. All participants provided written informed consent prior to testing and received monetary compensation for their time (10€/hour). Participants were randomly recruited from an online research participation system (Sona System), were naive to the purpose of the experiment, and reported no history of neurological, attentional, or psychiatric disorders. Research was conducted in accordance with the Declaration of Helsinki, and with ethical approval by the ethics committee of the International School for Advanced Studies (SISSA) (Protocol 10035-III/13). One participant was excluded from data analysis due to equipment failure (i.e., missing electroencephalography data). Thus, 21 participants were considered in the final sample of the study.

The sample size of the study was based on a previous study addressing magnitude integration with a similar paradigm, using dynamic stimuli (Togoli et al., 2021). To this aim, we performed a power analysis including the average effect size of the behavioural magnitude integration effect in that study (Cohen’s d = 1.31). Considering a two-tailed distribution and a power of 95%, we estimated a minimum required sample size of 10 subjects. However, since the present study also aimed to address the neural signature of magnitude integration with EEG, whose effect size is not known, we conservatively doubled such estimate aiming to test at least 20 participants.

### Stimuli

Visual stimuli were generated using the Psychophysics Toolbox (v.3; Kleiner M et al., 2007; Pelli, 1997) for Matlab (r2021b, The Mathworks, Inc.) and displayed on a 1092×1080 LCD monitor (120 Hz), encompassing a visual angle of 48×30 deg from a viewing distance of 57 cm. Similar to a previous study from our group (Togoli et al., 2021), dots were presented dynamically in a sequence of multiple, briefly-flashed (50-ms) dot arrays modulated in duration (i.e., the total duration of the sequence from the first to the last array) and in average numerosity (i.e., the average number of dots presented across the arrays composing a sequence). The numerosity of each individual array ranged from ±50% around the mean numerosity chosen for the specific trial, with the numerosity of each array computed ad-hoc in order for the overall sequence to result in a specific average numerosity. Dot positions in each array were computed to avoid dots overlapping with each other (minimum inter-dot distance = 2.5 times the radius of a single dot). The size of the dots and the size of the circular area encompassing them were systematically varied in a trial-by-trial fashion, following a procedure used in previous studies (DeWind et al., 2015; Park et al., 2016). The radius of the dots ranged from 6 to 10 pixels, while the field area of the stimulus (i.e., the virtual circular area where the dots were drawn) spanned from 200 to 400 pixel. Each array in the sequence had the same field area and the same dot size. Irrespective of the task condition (see below *Procedure*), all “probe” stimuli in the session were modulated in both duration and average numerosity, in a trial by trial fashion. The average numerosity of each probe stimulus could be either 15, 21, 30, 42, or 60 dots. The duration of each stimulus could be either 150, 200, 300, 450, 600 ms, which corresponded to the presentation of 3, 4, 6, 9, and 12 individual 50-ms arrays. The reference stimulus, presented at the beginning of the session and before the start of each block, had an average numerosity of 30 dots and a duration of 300 ms.

### Procedure

Each participant sat in front of a computer screen in a sound-attenuated room, and performed the duration and numerosity condition separately in two different sessions (2 hours each) taking place on different days in the same week. The order of the two conditions was counterbalanced across participants. Each condition involved a classification task of visual stimuli with the addition of EEG recording to measure both the behavioural effect of magnitude integration and the associated neural signature. In the classification task, participants had to make judgments about duration (duration task condition) or numerosity (numerosity task condition). Each of the five durations (150, 200, 300, 450, 600 ms) was combined with each of the five numerosities (15, 21, 30, 42, 60 dots) for a total of 25 possible combinations, randomly interleaved in a trial-by-trial fashion throughout each block of trials. At the beginning of the experiment, participants were shown a constant reference stimulus (30 dots, 300 ms) repeated ten times, and were asked to memorize it in order to be able to judge the probe stimuli in the main session. Once the block started, participants kept their gaze on a central fixation point, and a probe stimulus (i.e., a sequence of arrays defined by a given average numerosity and a given duration) was presented at the centre of the screen. After 600 ms from the offset of the probe stimulus, the fixation cross turned red signalling the end of the trial. The participant was then asked to judge, by pressing the left or the right arrow on the keyboard, whether the test duration or average number of dots was higher (right key) or lower (left key) compared to the memorized reference stimulus. Participants had 1200 ms to provide a response. If they did not press a button within this interval, the next trial started automatically (with an inter-trial interval, ITI, of 1100-1300 ms). Trials in which participants did not provide a response were excluded from data analysis. On average (± SD), participants missed 1.1% ± 1.2% and 0.9% ± 1.21% of the responses, respectively in the numerosity and duration task condition. After providing a response, the fixation point was flashed in green to confirm that the response was correctly registered, and the subsequent trial started automatically after a variable ITI of 1100 to 1300 ms. Participants received no feedback concerning the correctness of their response. At the beginning of each block after the first, the reference stimulus was repeated five times instead of ten. Note that the stimuli used in both conditions were similarly modulated across duration and numerosity, with the only difference being which dimension the participants were asked to attend and judge. Participants were asked to focus only on the task-relevant dimension (according to the condition), so that when judging duration, numerosity was irrelevant, and vice versa when judging numerosity. A total of 10 blocks of 100 trials each was collected in each condition, for a total of 1000 trials and 40 repetitions of each combination of numerosity and duration. Before the start of the session, subjects were familiarized with the task with a brief block of practice trials (∼ 10 trials).

### Behavioural Data Analysis

To assess the performance in the task and the potential biasing effects across magnitudes, we first computed the point of subjective equality (PSE) as a measure of accuracy, and the Weber fraction (WF) as a measure of precision. Individually for each subject and experimental condition, a psychometric function was fitted to the proportion of “more numerous” or “longer” responses as a function of the different levels of the task-relevant dimension. Moreover, to assess the magnitude integration effect, different psychometric fits were performed in the duration and numerosity task condition according to the different levels of the task-irrelevant, interfering dimension (numerosity in the duration task, duration in the numerosity task). From the cumulative Gaussian fit, the PSE was defined as the numerosity or duration level corresponding to chance level responses, which indicates a match in the perceived numerosity/duration of the probe with the memorized reference. The PSE represents the accuracy in the task and can be used to infer the perceived duration or numerosity of the probe stimuli as a function of the interfering magnitude dimension. The psychometric fitting was performed following the maximum likelihood method described by Watson, (1979). Additionally, to account for random errors and/or lapses of attention, we also applied a finger-error rate correction of 5% (Wichmann & Hill, 2001). What this correction does is to change the asymptotic levels of the probability range over which the cumulative Gaussian is fitted, with the rationale that a perfect proportion of 0 and 1 “test longer/more numerous” responses at the two extremes of the range may be prevented by random errors independent from the magnitude of the stimuli. With this correction, the cumulative Gaussian was thus fitted with a lower and upper boundary of 0.025 and 0.975, respectively. As a measure of precision in the task, we computed the Weber Fraction (WF), which is the ratio of the just noticeable difference (JND; i.e., difference in probe magnitude between 50% and 75% proportion of responses) and the PSE.

The JND, computed using the fitting procedure described above, is the minimum difference in duration or numerosity required to reliably discriminate test and reference stimuli. The JND value for each psychometric fit is thus computed from the slope of the curve (Watson, 1979), and scaled to reflect the difference in the magnitude of the test stimulus between the 50% and 75% levels of “probe more numerous/longer” responses. We chose the Weber fraction as measure of precision in the task as it considers the perceived magnitude of the stimuli, providing a more accurate index of precision if the variability of participants’ responses changes as a function of the stimulus magnitude (i.e., scalar variability; see for instance Cheyette & Piantadosi, 2020; Xu & Spelke, 2000), additionally, the WF provides a unit-less measure that can be compared across different tasks.

Finally, to obtain a better index of the magnitude modulation effect, a magnitude integration (MI) index was calculated. As shown in the formula below, the MI is based on the difference in PSE between the cases where test and reference stimuli have the same magnitude (PSE_baseline_), which was considered as a baseline measure of accuracy, and the cases in which the test had a higher or lower magnitude level compared to the reference (PSE_J_). This index was then turned into percentage and the sign was switched in order to facilitate its interpretation.

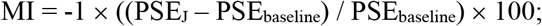

After the sign switch, a negative magnitude integration index thus indicates a relative underestimation of the probe stimulus compared to the baseline, while a positive index indicates an overestimation. The use of this magnitude integration index allowed us to directly compare the biases measured in different tasks, which would be otherwise very difficult to compare due to different measuring units of the PSE values (dots in the numerosity tasks, seconds in the duration task).

### Electrophysiological Data Acquisition and Pre-processing

While participants performed the numerosity or duration classification task, the EEG was recorded throughout the experimental procedure, using the Biosemi ActiveTwo system (at a sampling rate of 2048 Hz) and a 64-channel cap based on the 10-20 system layout. In addition, to monitor artifacts due to blinks or eye movements, the electro-oculogram (EOG) was measured via a channel attached below the left eye of the participant. The electrode offsets across channels were usually kept below 20 µV, but occasional values up to 30 µV were tolerated.

The pre-processing of EEG data was performed offline in Matlab (version R2021b), using the EEGLAB (Delorme & Makeig, 2004) and ERPlab (Lopez-Calderon & Luck, 2014) packages. During pre-processing, EEG signals were first re-sampled to a sampling rate of 1000 Hz, so that each data point corresponded to 1 ms. During the data pre-processing, each combination of duration and average numerosity was binned individually, for a total of 25 bins. The continuous EEG data was then segmented into epochs time-locked to the onset of each probe stimulus (i.e., the onset of the first array in each stimulus sequence), spanning from −300 ms to 1200 ms after stimulus onset. The pre-stimulus interval (−300:0 ms) was used for baseline correction, and the EEG signal was band-pass filtered with cut-offs at 0.1 and 40 Hz. To clean up the data from noise and artifacts, we first performed an independent component analysis (ICA) in order to remove identifiable artifacts such as eye movements and blinks. Additionally, we also employed a step-like artifact rejection procedure (amplitude threshold = 40 μV, window = 400 ms, step = 20 ms) to further remove any remaining large artifact from the EEG signal. This led to the exclusion of, on average, 2.9% ± 2.8% of the trials in the numerosity task condition, and 0.8% ± 0.9% of the trials in the duration task condition. Finally, the event-related potentials (ERPs) were computed by averaging EEG epochs within each bin. ERPs were then further low-pass filtered with a cut-off at 30 Hz.

### EEG Data Analysis

To assess the neural signature of magnitude integration, we performed a series of analyses on ERP data. First, we identified a set of a-priori channels of interest to assess the modulation of ERPs provided by duration and numerosity. Specifically, based on previous studies, we selected a series of six occipito-parietal channels, including O1, PO3, POz, PO4, O2, and Oz. Those channels were selected based on previous findings showing a signature of numerosity processing at occipito-parietal electrodes (Fornaciai et al., 2017; Park et al., 2016), and based on a recent study from our group showing a signature of distortions of perceived duration in a similar set of channels (Tonoyan et al., 2020). The rationale for this choice was to use the exact same set of channels when assessing the brain responses to both duration and numerosity, and their potential interaction, thus decreasing the degrees of freedom of the analysis. Across all the analyses and statistical tests, we thus always used the average of these six channels of interest.

First, to compare the time-course and strength of the effect of duration and numerosity on ERPs within and across the two task conditions, and assess any potential difference due to the task relevance of the different magnitudes, we computed the linear contrast of the ERPs sorted according to the different levels of either duration or numerosity. The linear contrast was computed according to the following weights: −2, −1, 0, 1, 2, corresponding to the five levels of either numerosity or duration. The contrast amplitude computed in this way was smoothed with a sliding-window average (50 ms, step = 10 ms). The contrast amplitude across each widow was tested with a series of one-sample t-tests against a null hypothesis of zero effect. The effect of numerosity and duration was them compared across task conditions using a series of paired-sample t-tests within each window. In both cases, multiple comparisons were controlled with a false discovery rate (FDR) procedure, with q = 0.05.

As the purpose of our study was to assess the interaction between numerosity and duration, we further assessed the extent to which the combination of duration and average numerosity modulated the brain responses to the stimuli. To do so, we sorted the ERPs according to the different combinations of the two magnitudes within each task condition, and employed a linear mixed-effect (LME) regression model to quantify the effect of the different magnitudes and their interaction, according to the following model:

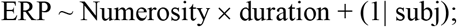

Where ERP refers to the amplitude of event-related potentials, numerosity refers to the different levels of average numerosity, and duration to the different durations of the stimuli. Subjects were added to this model as the random effect. The test was performed separately in the numerosity and duration task condition. This analysis was performed throughout the post-stimulus portion of the epochs (0:1200 ms), across a series of time windows spanning 50 ms each (step = 10 ms). The significance of each test across the different time windows was then collected and arranged in clusters of consecutive significant windows. To control for multiple comparisons, we first applied a cluster-size threshold of more than three consecutive time windows (i.e., we considered an effect as significant only when showing at least four consecutive significant windows). This cluster-size threshold was specifically chosen taking into account that individual time windows overlap with each other (i.e., having at least four time windows, the first and the last one overlaps by less than 50%). Moreover, we applied a cluster-based non-parametric test to assess the significance of each individual cluster corresponding to the effect of numerosity, duration, and their interaction. For each cluster of significant time-windows, we performed a simulation to assess how many times a similar cluster could be observed by chance. Namely, an LME regression test was performed at each window in a cluster, shuffling the ERP amplitude data corresponding to the different combinations of numerosity and duration. This test was repeated 1,000 times for each cluster, re-shuffling the ERP data each time, and the sum of t-values across the different windows was used as a cluster-level statistics. As a threshold to determine the significance of each cluster, we conservatively considered the minimum t-value computed in the actual cluster multiplied by the number of significant windows. To compute the cluster-level p-value, we then assessed the proportion of times the simulated clusters equalled or exceeded the threshold of the actual cluster.

Finally, based on the results of the LME analysis explained above, we selected four 100-ms latency windows corresponding to the significant interaction between duration and numerosity observed in either of the two conditions or both. Namely, we selected 0-100 ms, 360-460 ms, 800-900 ms, and 1050-1150 ms. Within each condition, we then computed the change in ERP amplitude as the difference in amplitude between each level of the interfering magnitude (duration in the numerosity task, numerosity in the duration task) and the middle magnitude level (300 ms/30 dots). We then performed a series of one-way repeated measures ANOVAs within each window, to assess whether the interfering magnitude induced a significant change in ERP amplitude. Additionally, we also assessed the relationship between the modulation of ERP amplitude calculated as a difference between each magnitude level and the middle value (as explained above) and the behavioural effect induced by the interfering dimension in the two task conditions. Namely, we assessed whether the strength of the behavioural effect, in terms of the magnitude integration index (MI), could predict the change in ERP amplitude induced by the modulation of the interfering magnitude. We then performed a linear mixed-effect model regression analysis at each latency window across the two task conditions, according to the following model:

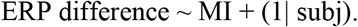

### Data availability

All the data generated during the experiments described in this manuscript can be found on Open Science Framework following this link: https://osf.io/vxdb7/. Experimental code will be made available by the corresponding author upon request.

## RESULTS

### Behavioural results

In the present study, participants performed either a numerosity or a duration classification task, whereby either the (average) numerosity or duration of each probe stimulus was judged according to a memorized reference. Each probe stimulus was composed of a series of briefly flashed (50-ms) dot arrays modulated in average numerosity (i.e., the average number of dots across the series of arrays; 15-60 dots) and duration (150-600 ms). Both dimensions were systematically modulated in the two tasks, so that the only difference across conditions was the dimension attended by participants and the task they performed. In the following sections, when mentioning numerosity we refer to the average numerosity across the different arrays composing each probe stimulus, unless otherwise specified.

First, we computed the point of subjective equality (PSE) as a measure of accuracy in the task and perceived magnitude of the stimuli, which is derived from a cumulative Gaussian (psychometric) fit to the proportion of “more numerous/longer” responses at different levels of the task-relevant magnitude. The PSE was computed separately for each level of the interfering dimension (duration in the numerosity task, numerosity in the duration task) in order to assess the behavioural effect of magnitude integration. The average PSEs across the two conditions is shown in Fig. 2A-B. As shown in the figure, the perceived magnitude of the stimuli varied noticeably as a function of the interfering magnitude, with a pattern showing a relative underestimation in the case of a lower interfering magnitude (i.e., higher PSE, suggesting that more dots or a longer duration is needed to perceptually match the reference) and overestimation in the case of a higher interfering magnitude.

**Figure 1.**
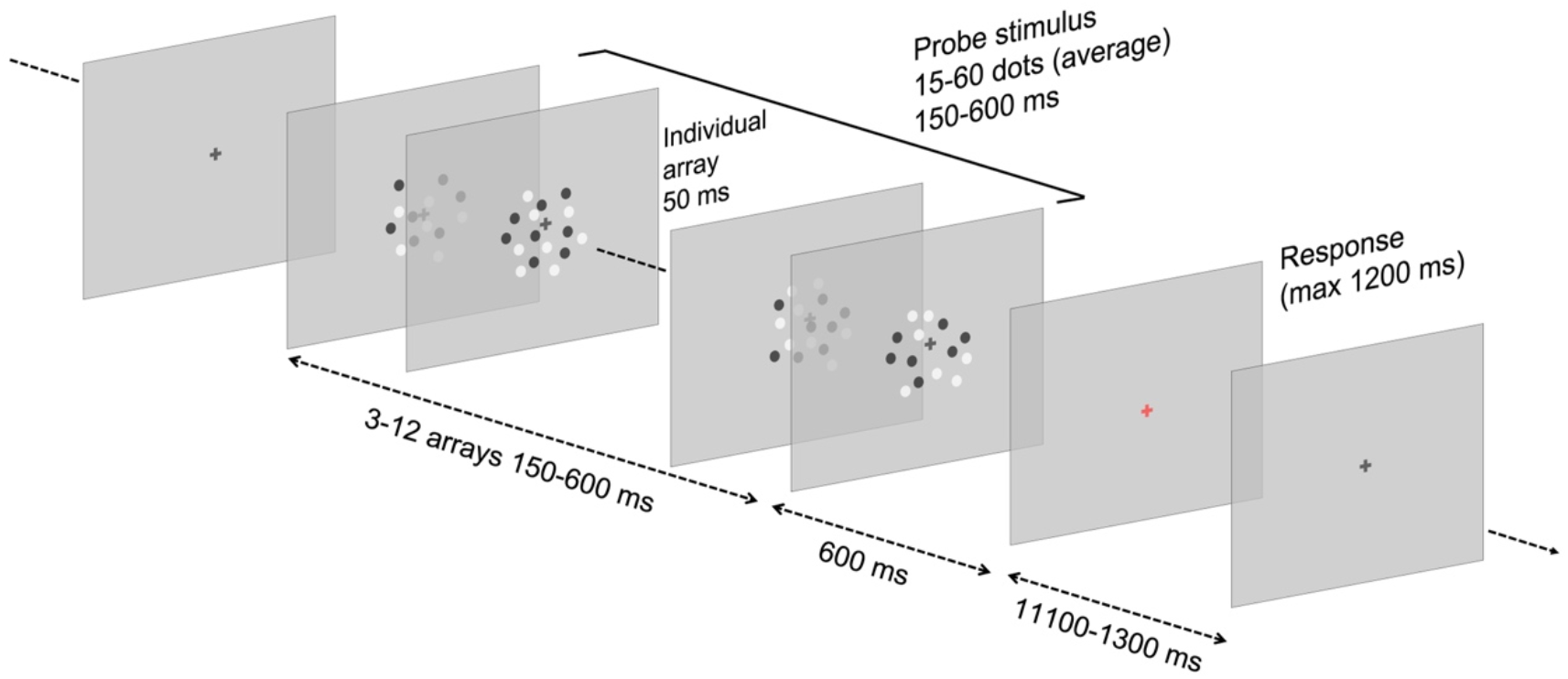
Example of the stimulation procedure. In the classification task, participants had to determine whether the probe stimulus was longer/shorter (duration condition) or more/less numerous (numerosity condition) compared to a memorized reference stimulus, presented at the beginning of the session and at the beginning of each block. Stimuli consisted in dynamic sequences of arrays varying in duration and in the average numerosity. Each array in the sequence included a cloud of white and black dots positioned within a circular area, presented for 50 ms. The number of arrays presented in each sequence depended on the duration selected in the trial and varied from 3 (150 ms) to 12 (600 ms). The offset of the stimulus sequence was followed by a 600 ms blank interval. After the interval, the fixation cross became red, signalling to the participant to provide a response by pressing the appropriate key on a standard keyboard. After providing a response, the fixation point turned green indicating that the response was recorded. The next trial started automatically after an inter-trial interval of 1100 to 1300 ms. Note that the stimuli are not depicted in scale.

**FIGURE 2.**
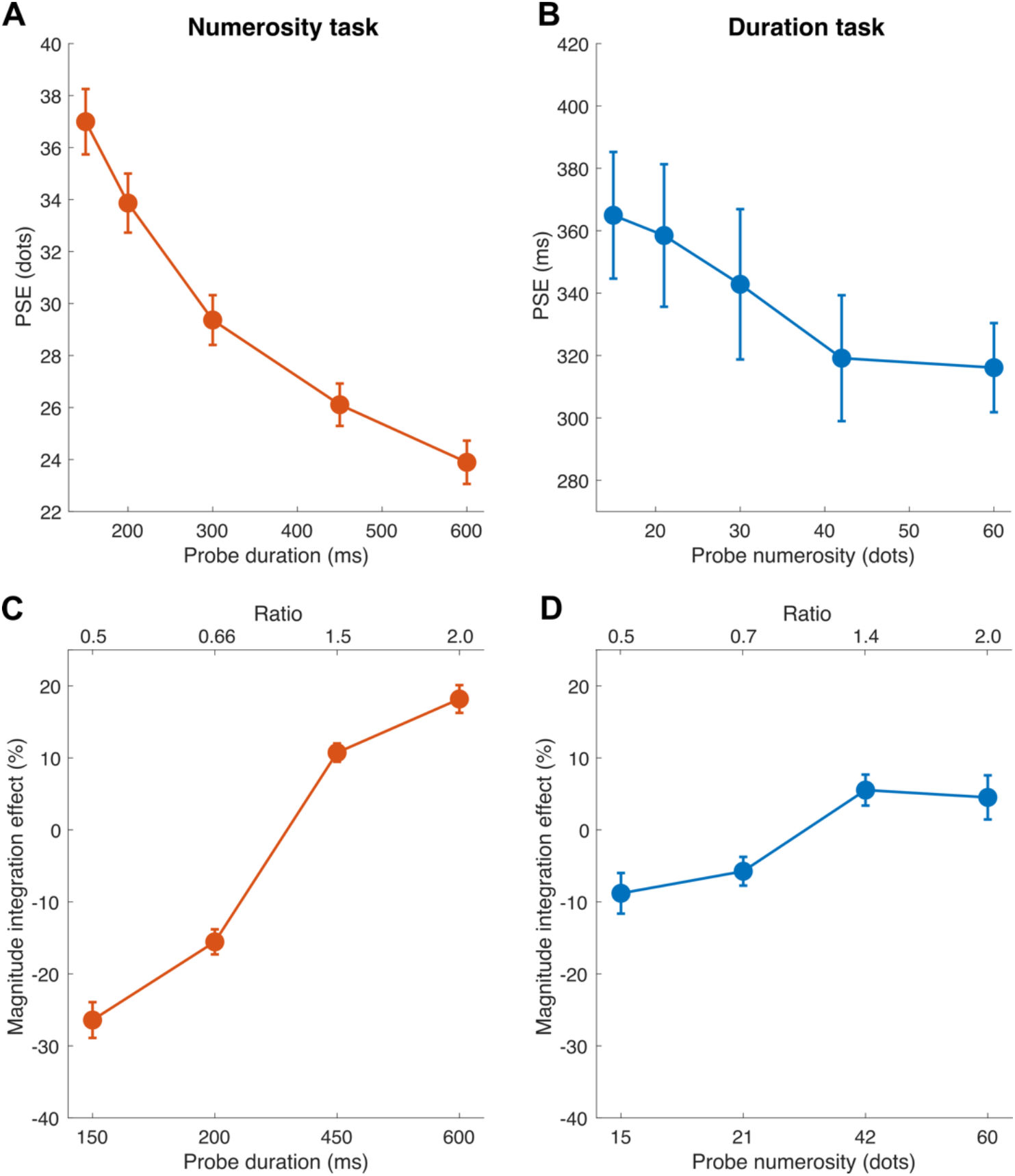
Behavioural results in terms of PSE and magnitude integration effect index. (A) Average PSEs observed in the numerosity task condition, as a function of the different durations of the stimuli (i.e., the interfering dimension). (B) Average PSEs observed in the duration task condition, as a function of the different numerosities of the stimuli. Note that a higher PSE means a relative increase in perceived magnitude compared to a lower PSE. Indeed, as in the classification task each stimulus was compared to a memorized reference, a higher PSE means that more dots or a longer duration is necessary to perceptually match the probe stimulus with the reference. (C) Magnitude integration effect index measured in the numerosity condition, as a function of the different duration levels. The magnitude integration effect index was computed as the difference in PSE between each level of the interfering dimension and the middle level (30 dots/300 ms), which was identical to the reference magnitude. In this case, we flipped the sign of the index in order facilitate its interpretation. Namely, a positive index shows an overestimation of the task-relevant magnitude, while a negative index shows an underestimation. (D) Magnitude integration effect index measured in the duration condition, as a function of the numerosity of the stimuli. Error bars are SEM.

To better assess the strength of magnitude integration and compare the effect measured in the two conditions, we computed a magnitude integration effect index (Fig. 2C-D) as the normalized difference in PSE between each level of the interfering dimension and the middle level of the range (which was equal to the reference). This allowed us to directly compare the effect obtained in the two different tasks. In the numerosity task (Fig. 2C), the estimated numerosity was biased by the duration of the stimuli: shorter durations compared to the reference (150 ms; 200 ms) led to an underestimation (MI = −26.43%; −15.65% respectively), while longer durations (450 ms; 600 ms) led to an overestimation (MI = 10.72%; 18.18% respectively). A similar, albeit weaker, pattern was found in the duration condition (Fig. 2D), where duration was underestimated (MI = −8.95%; −5.86%) for lower probe numerosities (15 and 21 dots respectively) and overestimated (MI = 5.65%; 4.45%) for higher ones (42 and 60 dots respectively). The pattern of magnitude integration effects across conditions was assessed with a linear mixed-effect (LME) regression model (MI ∼ ratio × task + (1|subj)), with “task” (numerosity or duration) and “ratio” (i.e., the ratio between the mid probe value and the other probe values) being the fixed effects, subjects the random effect, and the magnitude integration index (“MI”) as the dependent variable. The results of the LME model (R^2^ = 0.7) showed a significant effect of task (t = 4.91, p < 0.001) and ratio (t = −5,53, p < 0.001), but also a significant interaction between the two factors (t = −8.11, p < 0.001), which suggests an asymmetric influence across the two magnitudes. To better address the nature of this interaction, we first performed two individual LME regressions within each condition (MI ∼ ratio + (1|subj)). In the numerosity task (R^2^ = 0.8), the model showed a significant effect of ratio (t = −17.7, p < 0.001); similarly, also in the duration task (R^2^ = 0.6) we observed a significant effect (t = −6.2, p < 0.001). This shows that in both cases the interfering dimensions was effective in inducing a magnitude integration bias. Finally, we performed a series of paired t-tests to assess the strength of the effect at different levels of the interfering magnitude, in the numerosity vs. the duration task. Namely, the tests were performed across corresponding levels of ratio in the two conditions, considering a Bonferroni-corrected alpha of 0.0125. The results showed a significant difference across the magnitude integration index at most of the ratio levels (0.5: t(20) = 4.2, p < 0.001, Cohen’s d = 1.40; 0.66/0.7: t(20) = 3.22, p = 0.0043, d = 1.11; 2: t(20) = −3.9237, p < 0.001, d = 1.14), with the only exception of the ratio 1.4/1.5 (t(20) = −2.1329, p = 0.0455, d = 0.61). This further show that although the effect is significant in both directions, duration affected perceived numerosity to a larger extent compared to the effect of numerosity on duration.

Judging numerosity or duration usually entails different levels of difficulty. Indeed, at least in vision, judging duration is usually considered a more difficult task compared to a more salient dimension like numerosity, leading to a decreased precision of duration judgments compared to numerosity (Togoli et al., 2021). Although dynamic stimuli, whereby numerosity is defined as the average across several stimuli, have been shown to yield a similar precision of temporal and numerical judgments (Togoli et al., 2021), we further addressed this potential difference by assessing the Weber’s fraction (WF; computed as JND/PSE; see *Methods*) as a measure of precision in the task (Fig. 3A-B). As shown in Fig. 3, we observed a largely similar level of precision, ranging roughly from ∼0.19 to 0.21 in the numerosity task and from 0.22 to 0.24 in the duration task, with only a slightly lower precision in the case of duration judgments. We further performed a linear mixed-effect model regression (R^2^ = 0.73) on WFs (WF ∼ ratio × task + (1|subj)), with “task” and “ratio” as fixed effects and subject as random effect. The results showed no significant effect of ratio (t = 0.36, p = 0.71), task (t = −1.7, p = 0.08), and no significant interaction (t = −0.26, p = 0.8). This showed that WFs remained largely constant across the levels of the interfering dimensions, suggesting that an increase or decrease in PSE (bias) was accompanied by a proportional increase in the variability of responses (precision), in line with the Weber’s law (or scalar variability; Cheyette & Piantadosi, 2020; Xu & Spelke, 2000).

**FIGURE 3.**
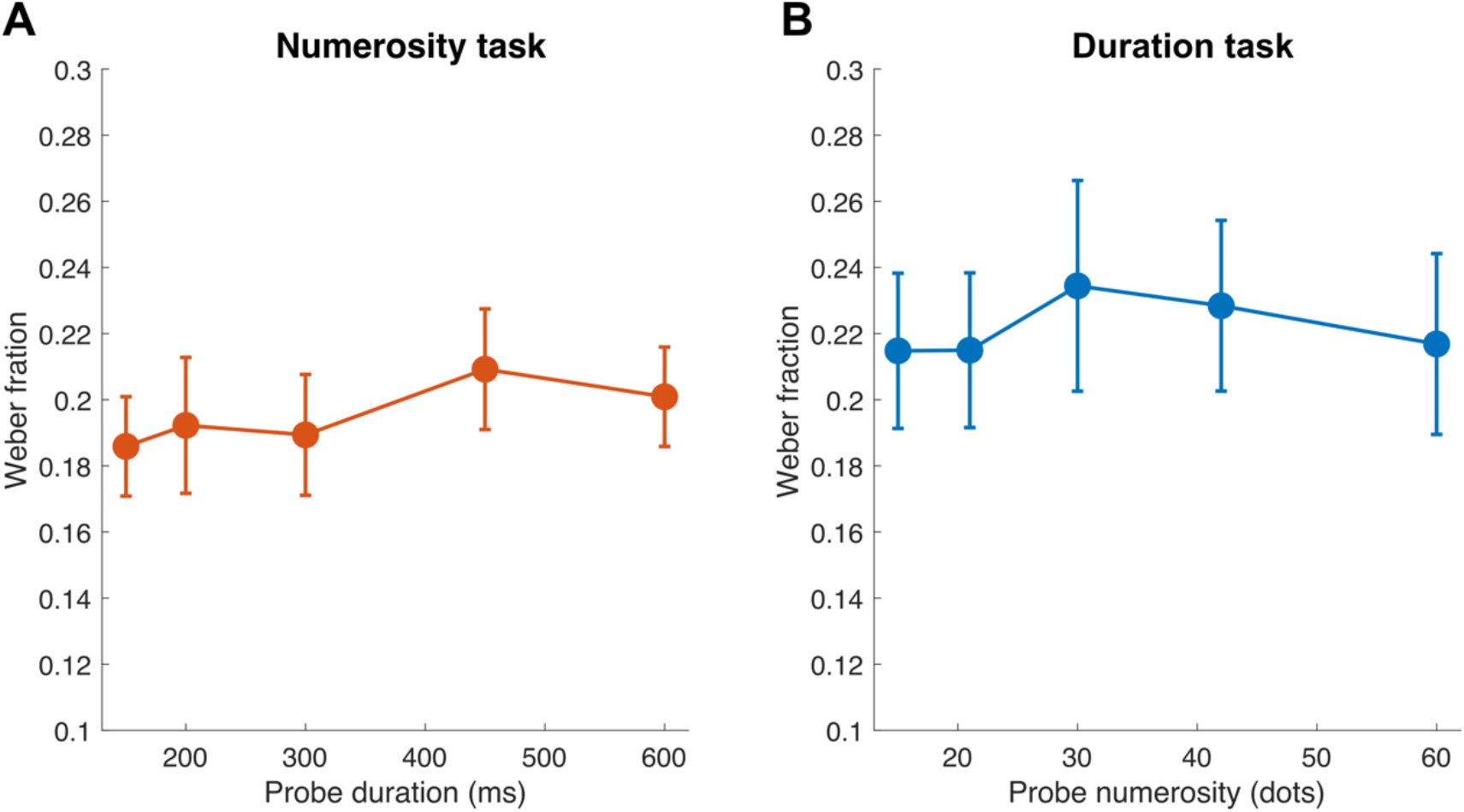
Weber’s fraction across the two conditions. (A) Weber’s fraction in the numerosity task condition, as a function of the different durations of the stimuli. (B) Weber’s fraction in the duration task condition, as a function of the different numerosities of the stimuli. Error bars are SEM.

### EEG results

Behavioural data showed systematic magnitude integration effects induced by duration on numerosity and by numerosity on duration. With EEG, we aimed to further address the neural signature of these effects, and assess the existence of a common magnitude processing stage mediating the interaction between numerosity and duration. The analysis of EEG data was performed by considering the average of a set of six occipito-parietal channels of interest in both the numerosity and duration task conditions (O1, PO3, POz, PO4, O2, and Oz), chosen according to previous results (Fornaciai et al., 2017; Park et al., 2016; Tonoyan et al., 2020).

Figure 4 shows the ERPs evoked by either different numerosities or different durations, separately for the two task conditions. In the case of the effect of numerosity, ERPs showed sensitivity to the numerical magnitude of the stimuli at around 150-200 ms, a latency potentially consistent with the N1 component, in both the numerosity (Fig. 4A) and duration (Fig. 4C) task. ERPs around this latency window appeared to show a more positive (or less negative) amplitude as a function of increasing numerosity. Sensitivity to numerosity seemed to emerge also at later latencies (500-700 ms after stimulus onset), again in both the numerosity and duration task condition. Duration, on the other hand, modulated ERPs at a later positive deflection, at around 300-400 ms in both task conditions (Fig. 4B and Fig. 4D), which may be consistent with the P3 component.

**FIGURE 4.**
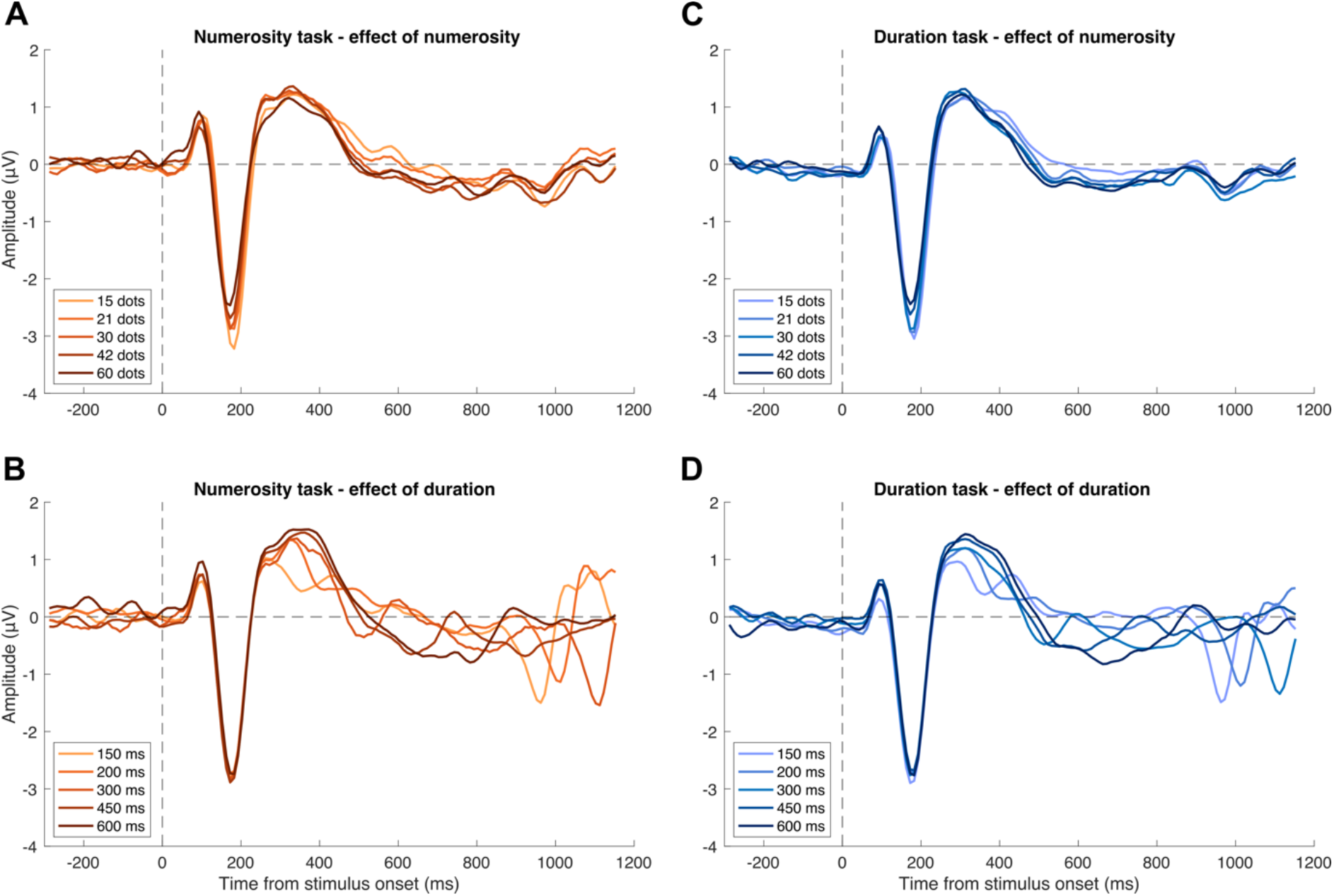
Event-related potentials evoked by different numerosities and durations. (A) Event-related potentials (ERPs) corresponding to different levels of average numerosity (i.e., effect of numerosity), measured in the numerosity task. (B) ERPs corresponding to different stimulus durations (i.e., effect of duration), in the numerosity task. (C) ERPs corresponding to different numerosities, in the duration task. (D) ERPs corresponding to different durations, in the duration task. ERPs were time-locked to the onset of the stimuli, which is marked by the vertical dashed line. The horizontal dashed line marks the zero in the amplitude scale.

First, we asked whether the manipulation of numerosity and duration could have a different impact on ERPs according to which stimulus dimension was attended by participants and which task they had to carry out. To do so, we computed the linear contrast across ERPs relative to all the levels of either numerosity and duration, within the numerosity and the duration task condition separately. If the modulation of visual evoked responses provided by different magnitudes depends on which dimension is task relevant and/or attended, then what we would expect is a difference in the extent to which the ERPs are modulated by the different magnitude levels. The different contrasts computed according to the different magnitudes and tasks are shown in Fig. 5.

**FIGURE 5.**
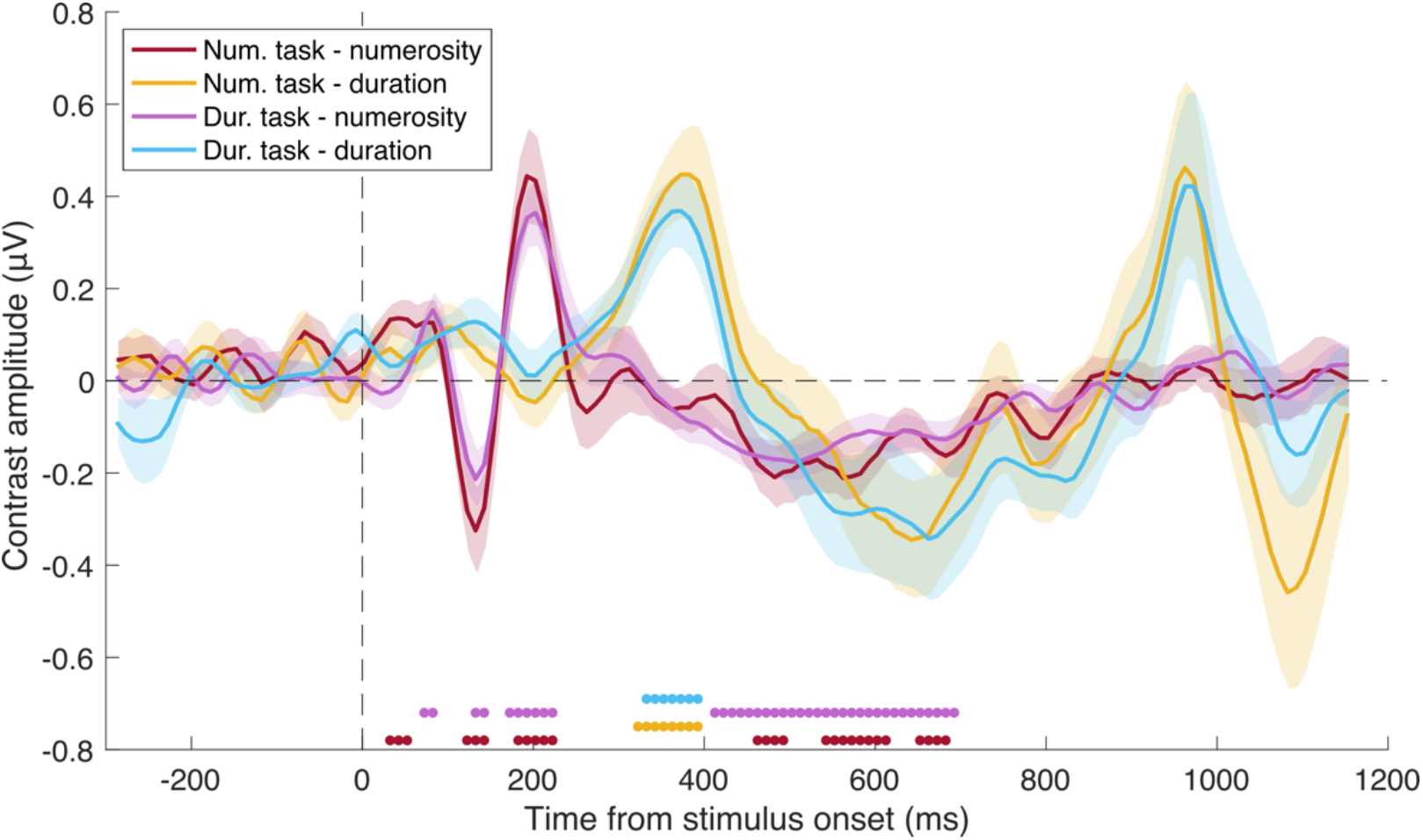
Contrast amplitude across the different dimensions and tasks. The linear contrast across the different magnitude levels of the two dimensions (contrast weights: [-2 −1 0 1 2]) was computed to assess whether visual evoked responses are differently modulated by either numerosity or duration according to which task participants were performing. Dots at the bottom of the plot show the significant time windows resulting from a series of one-sample t-tests against the null hypothesis of zero effect, with the same colour code as the main data. Shaded areas are SEM.

To address this point, we performed a series of one-sample t-tests against the null hypothesis of zero effect, corrected for multiple comparisons using a false discovery rate (FDR) procedure (q = 0.05). Regarding the modulation provided by numerosity on ERPs in the numerosity task, we observed three relatively early significant latency windows spanning 30-50 ms (t(20) ≥ 3.09, p ≤ 0.037), 120-140 ms (t(20) ≥ −3.08, p ≤ 0.037), and 180-220 ms (t(20) ≥ 3.32, p ≤ 0.027), followed by other three significant windows at later latencies: 460-490 ms (t(20) ≥ −2.95, p ≤ 0.042), 540-610 ms (t(20) ≥ −3.04, p ≤ 0.038), and 650-680 ms (t(20) ≥ −2.95, p ≤ 0.042). On the other hand, regarding the effect of numerosity on ERPs during the duration task, we similarly observed three significant early latency windows at around 70-80 ms (t(20) ≥ 3.88, p ≤ 0.007), 130-140 ms (t(20) ≥ −3.44, p ≤ 0.015), and 170-220 ms (t(20) ≥ 3.22, p ≤ 0.021), followed by a larger window at later latencies (410-730 ms; t(20) ≥ −2.73, p ≤ 0.046). Regarding the effect of duration on ERPs, in the numerosity task we observed a significant latency window at around 320-390 ms (t(20) ≥ 3.55, p ≤ 0.036). Similarly, in the duration task we observed a significant window at 330-390 ms (t(20) ≥ 3.54, p ≤ 0.042).

The crucial comparison to assess the possible effect of attention and task relevance however concerns the difference in the effect of numerosity and duration on the ERP contrast across the two tasks. With a series of (FDR-corrected) paired t-tests, we thus compared the effect of numerosity in the numerosity vs. the duration task, and the effect of duration in the numerosity vs. the duration task. The results of these tests however did not show any significant difference in the effect of the two different magnitude dimensions across the two tasks (all FDR-adjusted p-values ≥ 0.67). The modulation of visual evoked potentials provided by either duration or numerosity thus does not seem to depend on which dimension the participants are attending to or which task they are carrying out.

However, rather than the separate effect of numerosity and duration, we were more interested in the combined effect of these two magnitude dimensions. First, we thus assessed the extent to which the event-related potentials (ERPs) evoked by the stimuli in the two tasks are modulated by average numerosity, duration, or, crucially, the interaction between them. To do so, we sorted the ERPs according to each combination of average numerosity and duration, and performed a linear mixed-effect model regression (ERP ∼ average N × duration + (1|subj)). The ERPs corresponding to the combined levels of numerosity and duration (limited to the corresponding levels of the two ranges) are shown in Fig. 5. This analysis was performed by considering the average ERP amplitude across a series of 50-ms sliding windows (step = 10 ms) throughout the post-stimulus epoch (0-1200 ms). To determine the significance of the effects across the different time windows and control for multiple comparisons, we employed a cluster-based approach. Namely, first, we considered an effect as significant only when showing more than three consecutive significant time windows (four or more). Second, we performed a non-parametric test to assess the significance of each individual cluster observed in the regression analysis. Fig. 6 shows a subset of the different combinations of the two dimensions (i.e., the five “diagonal” combinations, out of a total of 25 combinations), and illustrates the different significant latency windows (i.e., clusters) observed in the regression analysis.

**FIGURE 6.**
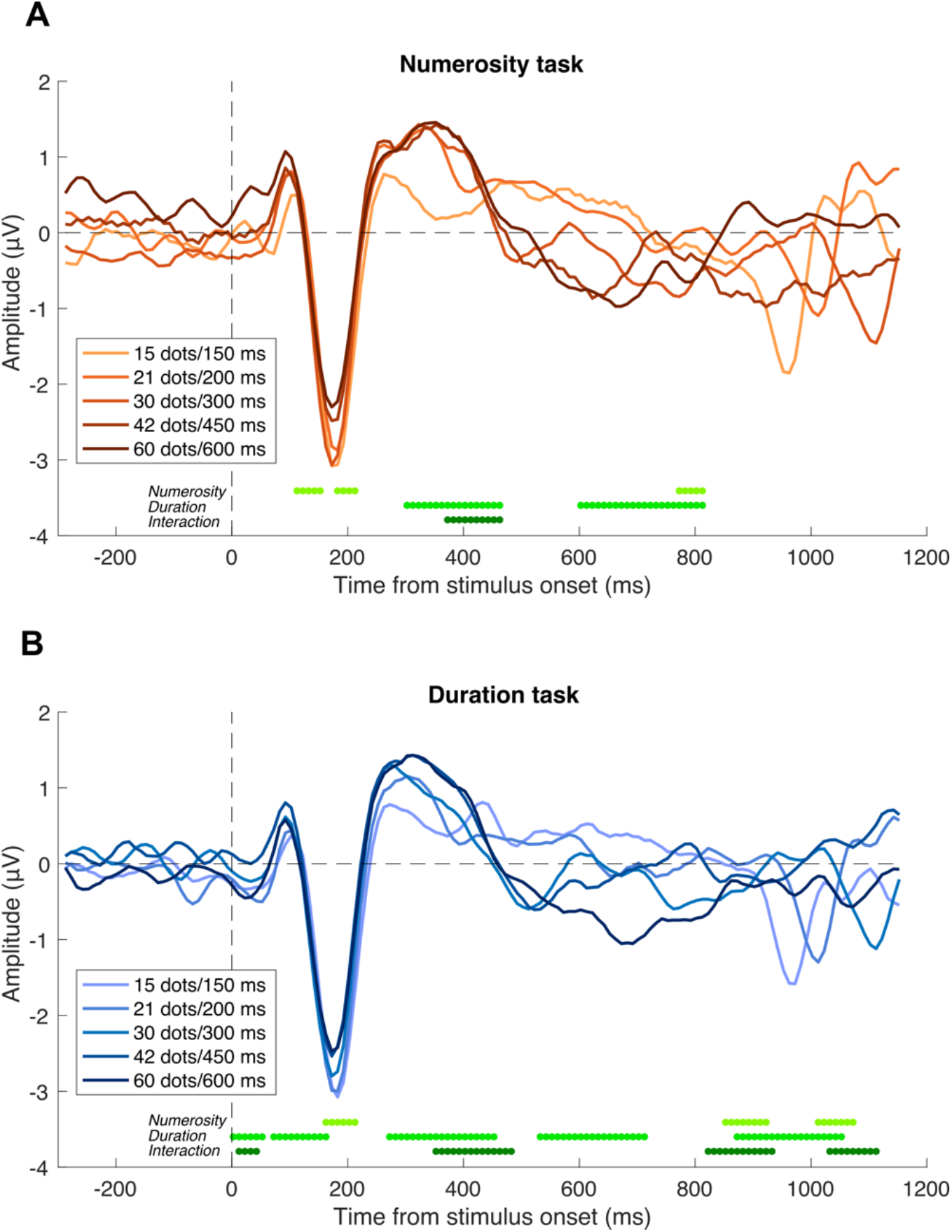
Event-related potentials (ERPs). ERPs time-locked to the onset of the stimuli (i.e., the onset of the first array in the stimulus sequence), plotted as a function of the congruent combination of increasing average numerosity and duration. Each wave represents the average of signals from six channels of interest at posterior scalp locations (O1, PO3, POz, PO4, O2, and Oz). (A) ERPs in the numerosity task condition, as a function of the combination of increasing average numerosity and increasing duration. (B) ERPs in the duration task condition. Lines in different shades of green at the bottom of each panel mark the timing of the different significant clusters observed in the linear mixed-effect model analysis, related to the effect of numerosity in modulating the amplitude of ERPs, the effect of duration, and the interaction between numerosity and duration (from top to bottom). Note that this figure only shows the ERPs related to the corresponding increasing levels of numerosity and duration; in the analysis, however, we considered the full set of 25 different combinations. ERPs shown here have been smoothed with a sliding window average (size = 50 ms, step = 10 ms) in line with the sliding widows used in the regression model.

As shown in Fig. 6, the modulation of different magnitude dimensions resulted in noticeable effects on the amplitude of ERPs, which appear to scale with the magnitude of the stimuli at several different latencies. For example, ERPs at around 100 ms, 200 ms, and 300 ms show an amplitude difference that orderly follow the increasing magnitude of the stimuli (i.e., more positive, or less negative amplitude for increasing magnitude). Signals at these different amplitudes may for instance be related to the P1, N1, and P2/P3 components, which in previous studies have been linked to numerosity and duration processing (Ernst et al., 2017; Fornaciai et al., 2017; Park et al., 2016; Temple & Posner, 1998).

In the numerosity task condition (Fig. 6A), the results of the linear mixed-effect model analysis showed a series of significant latency windows (i.e., significant clusters), relative to both the effect of numerosity, duration, and their interaction. Regarding the influence of numerosity, we observed three clusters of significant effect, at 110-150 ms (average R^2^ = 0.80 ± 0.025; α ranging from −0.37 to −0.73, t-values ranging −2.18 to −4.04, p-values ranging from 0.015 to 0.0006; cluster-based p-value < 0.001), 180-210 ms (R^2^ = 0.88 ± 0.004; α = 0.44 to 0.60, t = 2.57 to 3.52, p = 0.037 to 0.014; cluster-based p-value < 0.001), and 690-810 ms (R^2^ = 0.39 ± 0.018; α = −0.51 to −0.61, t = −1.62 to −2.46, p = 0.037 to 0.014; cluster-based p-value < 0.001). Regarding the effect of duration, we observed two clusters of significant effect, at around 300-460 ms (R^2^ = 0.78 ± 0.061; α = 0.33 to 1.14, t = 2.06 to 6.34, p = 0.04 to <0.0001; cluster-based p-value < 0.001) and 600-840 ms (R^2^ = 0.34 ± 0.059; α = −0.45 to −0.72, t = −1.97 to −3.52, p = 0.049 to 0.0005; cluster-based p-value < 0.001). Finally, but more importantly, we observed a significant interaction between numerosity and duration at about 360-460 ms (R^2^ = 0.75 ± 0.068; α = −0.31 to −0.48, t = −2.08 to −2.87, p = 0.04 to 0.004; cluster-based p-value < 0.001).

In the duration task condition (Fig. 6B), we observed a larger number of significant clusters, showing modulations of ERP amplitude provided by numerosity, duration, and their interaction. First, regarding the effect of numerosity on ERPs, we observed a total of three significant clusters: 160-210 ms (average R^2^ (± SD) = 0.88 ± 0.012; α = 0.56 to 0.70, t = 2.19 to 4.75, p = 0.03 to <0.0001; cluster-based p-value < 0.001), 850-920 ms (R^2^ = 0.23 ± 0.057; α = 0.46 to 0.55, t = 1.97 to 2.39, p = 0.049 to 0.018; cluster-based p-value < 0.001), and 990-1070 ms (R^2^ = 0.02 ± 0.007; α = 0.57 to 0.68, t = 1.97 to 2.21, p = 0.049 to 0.027; cluster-based p-value < 0.001). On the other hand, in terms of the effect of duration on the ERP amplitude, we observed five clusters of significant effects. The clusters spanned 0-50 ms (R^2^ = 0.52 ± 0.042; α = 0.31 to 0.47, t = 2.12 to 3.17, p = 0.034 to 0.001; cluster-based p-value < 0.001), 70-160 ms (R^2^ = 0.77 ± 0.101; α = 0.28 to 0.34, t = 2.00 to 2.38, p = 0.044 to 0.02; cluster-based p-value < 0.001), 270-450 ms (R^2^ = 0.80 ± 0.063; α = 0.28 to 0.97, t = 1.97 to 6.143, p = 0.049 to <0.0001; cluster-based p-value < 0.001), 530-710 ms (R^2^ = 0.44 ± 0.059; α = −0.37 to −0.66, t = −2.10 to −3.49, p = 0.036 to < 0.001; cluster-based p-value < 0.001), and 870-1050 ms (R^2^ = 0.09 ± 0.088; α = 0.46 to 1.11, t = 2.21 to 3.50, p = 0.027 to < 0.001; cluster-based p-value < 0.001). Finally, we also observed a series of clusters showing a significant effect of the interaction between numerosity and duration in modulating ERPs. Namely, a significant interaction was observed at 10-40 ms (R^2^ = 0.52 ± 0.026; α = −0.29 to −0.40, t = −2.14 to −2.95, p = 0.032 to 0.003; cluster-based p-value < 0.001), 350-480 ms (R^2^ = 0.72 ± 0.092; α = −0.31 to −0.45, t = −2.26 to −3.14, p = 0.032 to 0.001; cluster-based p-value < 0.001), which overlaps with the interaction observed in the numerosity task, 820-930 ms (R^2^ = 0.24 ± 0.073; α = −0.40 to −0.62, t = −2.10 to −2.97, p = 0.036 to 0.003; cluster-based p-value < 0.001), and 1030-1110 ms (R^2^ = 0.01 ± 0.002; α = −0.54 to −0.64, t = −2.00 to −2.33, p = 0.045 to 0.020; cluster-based p-value < 0.001).

Overall, this analysis showed that the modulation of different magnitude dimensions and, importantly their interaction, had a strong effect on ERPs at several different latencies after stimulus onset, throughout the entire stimulus epoch. Interestingly, we found a common latency window showing a significant interaction between numerosity and duration, with a similar timing across both task conditions. This latency window may thus qualify as a potential signature of a shared processing stage of different magnitude dimensions. Observing an interaction between duration and numerosity is however not sufficient to conclude that a given processing stage supports magnitude integration, as the interaction itself does not tell us whether the evoked activity is consistently modulated by the interfering dimension in a congruent fashion. A genuine correlate of magnitude integration should indeed (1) show activity parametrically modulated by the interfering magnitude, and (2) a relationship with the magnitude integration bias measured behaviourally.

We thus identified a series of target latency windows based on the interaction effects observed in the regression model (i.e., the windows showing a significant interaction in either of the two tasks or both), and performed further analyses to address the pattern of activity across them and assess the relationship with the behavioural effect. Namely, we selected four 100-ms latency windows, spanning 0-100 ms, 360-460 ms, 800-900 ms, and 1050-1150 ms. Within each window, we computed a measure of “ERP difference” based on the contrast between each level of the interfering magnitude (150, 200, 450, 600 ms in the numerosity task, 15, 21, 42, 60 dots in the duration task) and the middle level of the range (300 ms/30 dots) – similarly to what we did to compute the behavioural magnitude integration effect index. The pattern of ERP difference across the different windows and tasks is shown in Fig. 7.

**FIGURE 7.**
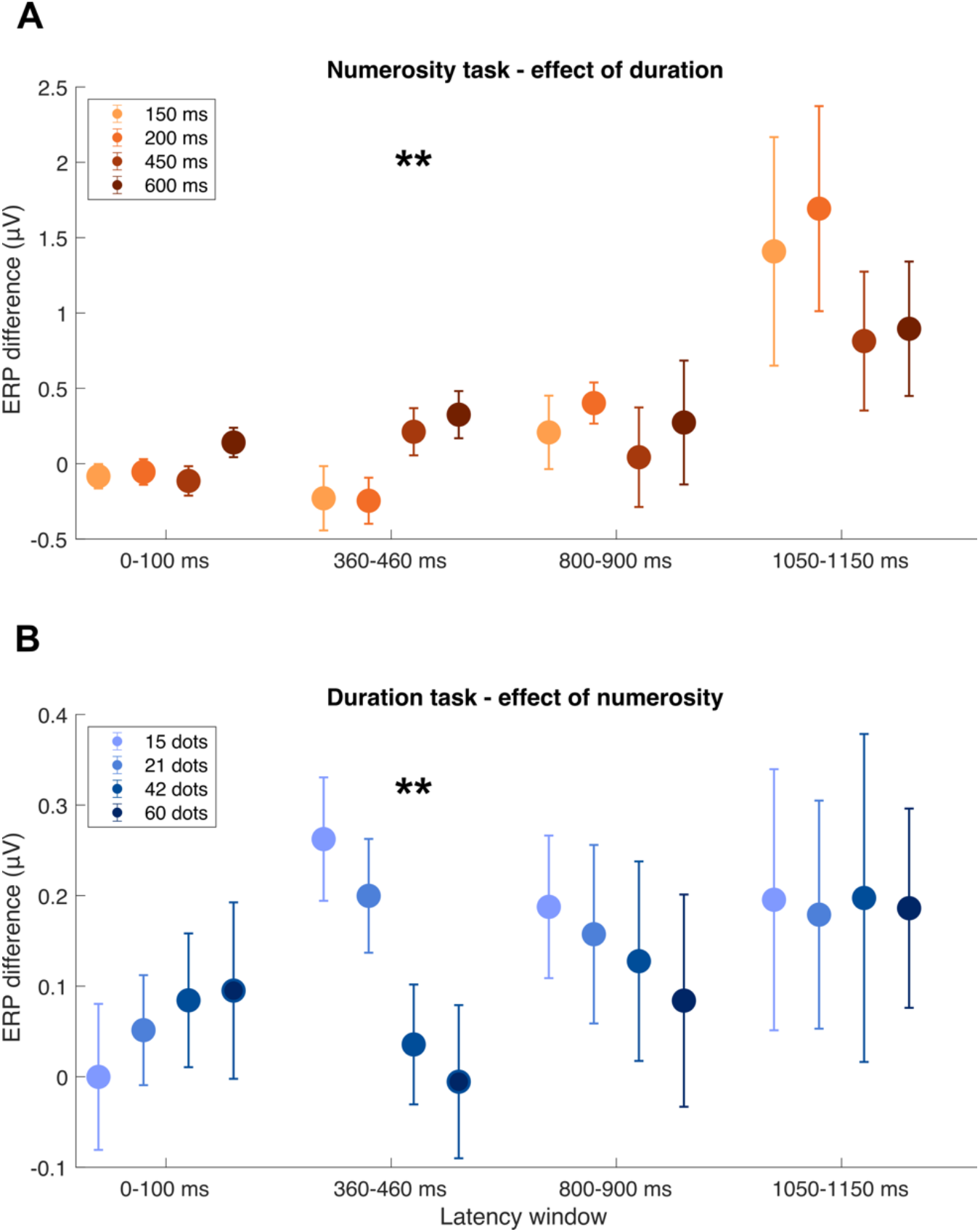
Differences in ERP amplitude as a function of the interfering magnitude within different latency windows. (A) ERP difference across the four latency windows in the numerosity task condition, indexing the effect of duration on the amplitude of ERPs. (B) ERP difference in the duration task condition, indexing the effect of numerosity on the amplitude of ERPs. Error bars are SEM. Stars refer to the significance of one-way repeated measures ANOVA performed within each latency window. ** p < 0.01.

First, we assessed how the ERP amplitude within these windows is modulated by duration in the numerosity task condition, and by numerosity in the duration task condition, in terms of the ERP difference measure. We thus performed a series of one-way repeated measures ANOVAs within each window, with factor either duration or numerosity. In the numerosity task condition (Fig. 7A), we found that activity within the 360-460 ms latency window showed a significant main effect of duration (F(3,60) = 5.19, p = 0.003, η_p_^2^ = 0.21). Although not completely parametric, activity in this latency window indeed showed a consistent modulation provided by the interfering dimension, with a positive or negative change in ERP amplitude as a function of whether the duration was longer or shorter than the middle value. None of the other latency windows showed significant effects (0-100 ms: F(3,60) = 1.97, p = 0.128, η_p_^2^ = 0.09; 800-900 ms: F(3,60) = 0.50, p = 0.681, η_p_^2^ = 0.02; 1050-1110 ms: F(3,60) = 2.44, p = 0.073, η_p_^2^ = 0.11). Similarly, in the duration task condition (Fig. 7B) we observed a significant effect of numerosity on the ERP amplitude in the 360-460 ms window (F(3,60) = 5.43, p = 0.002, η_p_^2^ = 0.21). In this case, the direction of the effect was reversed, with higher numerosity decreasing the ERP amplitude (i.e., negative ERP difference) compared to the middle value, but again in a way that is consistent with the different magnitude levels, in a parametric fashion. Also the 0-100 ms and 800-900 ms latency windows showed seemingly consistent effects, but not strong enough to reach significance (0-100 ms: F(3,60) = 0.54, p = 0.65, η_p_^2^ = 0.03; 800-900 ms: F(3,60) = 0.42, p = 0.736, η_p_^2^ = 0.02; 1050-1110 ms: F(3,60) = 0.009, p = 0.99, η_p_^2^ = 0.001). This analysis suggests that the pattern of ERP activity within the 360-460 ms latency window is modulated by the interfering dimension in both task conditions, while activity across the other windows does not show a clear pattern.

Then, we went on and addressed the potential link between ERP activity within these four latency windows and the magnitude integration effect measured behaviourally. To do so, we employed a linear mixed-effect model regression on the average activity across the window, with the magnitude integration effect index as fixed effect factor, and subjects as random effect (ERP difference ∼ MI + (1|subj)). In the numerosity task condition (Fig. 7A), we observed again a significant effect within the 360-460 ms latency window (R^2^ = 0.43, α = −1.10, t = −2.79, p = 0.006). No other significant effect was observed across the other windows (0-100 ms: R^2^ = 0.13, α = −0.24, t = −1.04, p = 0.30; 800-900 ms: R^2^ = 0.53, α = 0.33, t = 0.56, p = 0.57; 1050-1150 ms: R^2^ = 0.83, α = 1.28, t = 1.64, p = 0.10). These results show that, limited to the 360-460 ms latency window, the modulation of ERPs can be significantly predicted based on the strength of the behavioural magnitude integration effect. In the duration task condition (Fig. 6B), we observed two significant latency windows: again, the 360-460 ms latency window (R^2^ = 0.37, α = 0.76, t = 2.61, p = 0.01), and the 800-900 ms window (R^2^ = 0.62, α = 0.92, t = 2.63, p = 0.01). No significant effect was instead observed at the 0-100 ms (R^2^ = 0.49, α = −0.33, t = −1.10, p = 0.27) and 1050-1150 ms (R^2^ = 0.68, α = 0.77, t = 1.68, p = 0.09) window.

## DISCUSSION

In the present study we investigated the neural signature of magnitude integration between numerosity and duration, to address the nature of this phenomenon and its possible neural mechanisms. To do so, we employed a numerosity and a duration classification task of dynamic dot-array stimuli composed of a series of briefly-flashed arrays with varying numerosity. In both tasks, the stimuli were systematically modulated in the dimension of average numerosity (the average number of dots across all the arrays composing a stimulus sequence), and in duration. Importantly, the stimuli were identical in the two task conditions, and the only difference across them was which dimension the participants were asked to attend and judge.

First, our behavioural results show a robust pattern of magnitude integration effects in both task conditions, with perceived numerosity affected by duration in a congruent fashion (i.e., the longer the stimulus, the higher the perceived numerosity) and perceived duration affected by numerosity (i.e., the more numerous the stimulus, the longer the perceived duration). Although the effect was significant in both directions, we observed an asymmetry: the influence of duration on numerosity was stronger compared to the influence of numerosity on duration. Asymmetric effects in magnitude integration are common in the literature, but usually involve duration being more strongly affected by non-temporal dimensions (e.g., Casasanto & Boroditsky, 2008; Dormal et al., 2006; Togoli et al., 2021; but see Cai & Connell, 2015 for an opposite asymmetry in the tactile modality). Symmetric effects across duration and numerosity have been observed by using either very short stimulus durations (i.e., ∼50-100 ms; Javadi & Aichelburg, 2012) or dynamic stimuli whereby numerosity is conveyed as an average across several individual arrays (Togoli et al., 2021). This has led to the proposal that asymmetric effects across duration and numerosity result from a different processing time-course of these two dimensions, with the fastest one to be represented (i.e., numerosity; see Fornaciai et al., 2017) being able to more strongly interfere with a slowly accumulating dimension (i.e., duration), but not vice versa (Togoli et al., 2021); see also Lambrechts et al., 2013). When using average numerosity, instead, the processing time-course of numerosity becomes more similar to duration, as it involves tracking information over time. One difference compared to our previous study (Togoli et al., 2021) is however the task used and how the stimuli are modulated in the two dimensions. Indeed, in Togoli et al., (2021) we employed a discrimination task involving the comparisons of two stimuli: a reference modulated in the interfering dimensions and a probe modulated in the task-relevant dimension. Here, instead, participants compared each probe stimulus, which was concurrently modulated in both the task-relevant and interfering dimension, with a memorised reference. In this case, differences in how the reference is memorised and used to perform the task might influence the strength of the effect, as the effect itself is based on the probe interfering dimension being perceived as higher or lower than the reference. Our results however do not allow to draw a strong conclusion on the nature of this asymmetry, which thus remains an open point. In any case, although this asymmetry is potentially interesting and worth further investigation, the crucial aspect of our results is that they show significant magnitude integration biases both from numerosity to duration and from duration to numerosity.

The presence of mutual biases across duration and numerosity in our paradigm is indeed an essential precondition in order to be able to address the neural signature of magnitude integration and the existence of a common neural processing stage mediating it. To this aim, the EEG was recorded throughout the two task conditions, in order to assess the brain responses to the different combinations of the two dimensions. In our hypothesis, the conditions that a neural signature should meet to be considered as evidence for a shared processing stage mediating magnitude integration are: (1) brain activity should be modulated by the interaction of numerosity and duration, and (2) the pattern of activity at such a stage should be consistently (i.e., parametrically) modulated by the interfering dimension; (3) such a processing stage should be independent from attention and task-related factors (i.e., task relevance of different dimensions); (4) finally, the pattern of brain activity at that stage should be related to the magnitude integration bias measured behaviourally.

In our analysis, we identified a unique latency window meeting all these criteria, and thus supporting the existence of a shared processing stage where numerosity and duration are integrated giving rise to the behavioural bias. First, assessing the impact of numerosity and duration on visual evoked potentials showed several latency windows systematically modulated by either one or the other dimension (see Fig. 4-6). Across the two task conditions, the most prominent signature of sensitivity to numerosity was observed in a negative ERP deflection at around 150-200 ms, which is consistent with the timing of the N1 ERP component. Previous studies on numerosity perception typically showed numerosity-sensitive ERPs around similar latencies (∼200 ms after stimulus onset) and at similar posterior occipito-parietal channels, although usually with a positive deflection (P2p component; Fornaciai et al., 2017; Park et al., 2016; Temple & Posner, 1998; but see also Xuan et al., 2009). However, previous studies investigating the neural effects of numerosity usually employed static dot-array stimuli, making it difficult to directly compare the results. Additional significant effects of numerosity were observed at much later latencies (∼800 ms and 900-1000 ms, respectively in the numerosity and duration task condition), which are unlikely to be related to perceptual numerosity processing, and could instead be linked to the more cognitive or behaviour-related aspects of the task. Note however that the latest latency window observed in the duration task showed a very low R^2^ (0.02), making it an unreliable result.

On the other hand, the modulation of ERPs provided by duration showed a much more widespread signature at several different latencies. Across the two tasks, two common latency windows showed a significant effect, at around 350-400 ms and 600-700 ms. The first of these two windows may be consistent with the P3 component, which earlier findings by Ernst et al., 2017 suggested to be potentially linked to perceived time. Interestingly, in the study by Ernst et al., (2017), the P3 amplitude reflected the overestimation of duration induced by oddball stimuli rather than duration per se. In the present results, part of the latency window showing a significant effect of duration overlapped with a slightly later window showing a significant interaction between numerosity and duration, suggesting that similarly to Ernst et al., (2017) this latency reflects a distortion in perceived time. However, considering the interaction, activity in this latency window seems to be also modulated, at least in part, by duration per se (see below for more discussion about the interaction). The later latency window spanning from ∼600-800 ms (numerosity task) or ∼500-700 ms (duration task), which is associated with a negative deflection of ERPs, might instead be consistent with the contingent negative variation component (CNV). For example, in Zhang et al., (2021), the timing of the CNV was roughly similar compared to the present study. However, the involvement of the CNV with perceptual timing is debated, and several studies suggest that it is related to decision-making (see Kononowicz & Penney, 2016 for a review), and it is affected by memory and temporal context (Wiener & Thompson, 2015; Zhang et al., 2021). Importantly, one key feature of the CNV is its relationship with the specific decision criterion on the experiment (Macar & Vidal, 2003; Pouthas et al., 2000), showing a deflection closer to the decision boundary allowing to classify or discriminate the stimuli (i.e., for instance the duration used as reference). The timing of the effect of duration shown in our results is thus not consistent with the CNV (i.e., our reference was 300 ms). Additionally, the CNV usually peaks at fronto-central electrodes, while our analysis only involved posterior occipito-parietal channels. Interestingly, in the duration task only, we also observed an early effect of duration, with a significant window at ∼0-50 ms, which overlaps with a significant interaction with numerosity, and a second one at ∼50-150 ms. This latter window is consistent with the timing of the P1 component, and might for instance be related to the initial stages of temporal accumulation that will influence perceived duration, similarly to what has been suggested in auditory time perception by Bendixen et al., (2005). Finally, we also observed a later latency window at around ∼900-1050 ms, but, since the regression models showed an extremely low R^2^ (0.09), it is difficult to consider this a reliable result.

Besides the individual effects of numerosity and duration in modulating the brain responses evoked by the stimuli, we were particularly interested in the interaction between numerosity and duration, in order to assess the existence of a shared magnitude processing stage. Across the two conditions, we observed a specific latency window showing a significant interaction between the two dimensions, at around 360-460 ms in the numerosity task, and 350-480 ms in the duration task. Such an effect emerging at overlapping latencies suggests a specific processing stage independent from the task performed by participants. Besides this latency window, we also observed a few more windows showing a significant interaction, but only in the duration condition. Namely, a very early one (∼0-50 ms), and two later windows at around ∼800-1100 ms. While it is difficult to interpret the nature of these effects, we can speculate that they may be related to either the earliest stages of temporal accumulation which may be jointly modulated by duration and numerosity (0-50 ms; see Fornaciai et al., 2017 for a signature of numerosity processing at very early latencies), or reflect response-related or decisional processes (800-930 ms). We also observed an even later interaction at 1030-1110 ms, but the extremely low R^2^ in this window (0.01) suggests again that this is not a reliable result.

To further address these latency windows according to our criteria for a common magnitude processing stage, we assessed the impact of duration and numerosity within these windows (Fig. 6). Our results demonstrate that only the 360-460 ms latency window shows a meaningful a pattern of activity entailing a significant effect of the interfering magnitude and a significant relationship with the behavioural magnitude integration effect. Specifically, in both the numerosity and duration task, the effect of magnitude on ERPs seems to scale according to the magnitude of the interfering dimensions, or at least show a congruent pattern according to whether the magnitude was lower or higher compared to the middle level of the range. This overlapping timing across the two tasks and the fact that only this window shows a meaningful and significant effect of the interfering magnitude suggests that this is a key stage potentially related to magnitude integration. The significant relationship with the behavioural effect of magnitude integration further supports this interpretation. Namely, the strength of the ERP modulation provided by the interfering dimension is significantly related to the strength of the behavioural effect. Additionally, we also observed a significant relationship between ERPs and behavioural effects at a later window (800-900 ms). This effect is however specific for the duration task condition, and the pattern of ERP difference at that stage does not show a significant effect of the interfering magnitude, but just a trend. This stage could thus be related to some additional decision- or response-related processing occurring specifically in the duration task, and its unlikely to represent a genuine magnitude integration effect.

A processing stage taking place at around 360-460 ms after stimulus onset seems thus to reflect the integration of different magnitude dimensions leading to the behavioural bias. However, what is the nature of such processing stage? Is it related to the perceptual processing of the stimuli, or to more cognitive processes such as memory encoding or decision-making? When it comes to interpreting the nature of such a magnitude processing stage, we first reasoned that the timing of the latency window where it occurs might provide some information. However, as it is difficult to draw a clear line between the timing of perceptual and cognitive processes, any interpretation remains speculative. In our hypothesis, we first defined as “perceptual” processing in broad terms the time-course of evoked response within our duration range (i.e., 0-600 ms). Due to the dynamic nature of the magnitudes investigated in our study, we thus expect that brain processes during this time would concern the accumulation of numerical and temporal information and the updating of perceptual representations with incoming information. Considering the duration range used, the critical latency window for a perceptual magnitude integration can be narrowed down to after the average offset of the stimuli, which is 300-600 ms. A “final” representation of average numerosity and duration can indeed occur only when the entire stimulus sequence has fully unfolded. Since all the durations were included in this analysis, we thus expected the critical time for perceptual representation and magnitude integration to be shortly after the middle of our duration range. This is indeed in line with what we observed, as a significant interaction between numerosity and duration was observed in the 360-460 ms latency window. An alternative interpretation of this specific latency however involves the possibility of decisional processes crossing an important decision boundary – i.e., the duration of the stimuli moving from shorter (< 300 ms) to longer (> 300 ms) than the reference. Such a decisional process might in turn be involved in determining the pattern of magnitude integration. Thus, the timing of the effect alone is not sufficient to draw a strong conclusion concerning the nature of the effect and the integration mechanism.

To better disentangle the role of perceptual versus decisional or mnemonic processes, the extent to which attention and task-related factors affect the signature of different magnitudes might provide further information. Our reasoning in this context is that, although it is known that irrelevant stimulus dimensions can be encoded in visual working memory (e.g., Gao et al., 2016), the encoding of irrelevant information is expected to be weaker (Bocincova & Johnson, 2019). This, in our experiment, should thus lead to a decreased impact of numerosity and duration modulation on ERPs in the different tasks, when they are irrelevant to current task at hand (i.e., weaker effect of numerosity in the duration task compared to the numerosity task, and vice versa for duration). In terms of perceptual processing, instead, no difference would be expected in the effect of different stimulus dimensions according to the task. Our results, concerning the linear contrast of ERPs evoked by different levels of numerosity and duration (Fig. 5), show a very similar pattern of modulation across the two tasks, and no significant difference. Moreover, the involvement of decisional processes should be related to a similar asymmetry across different tasks, as the two dimensions are expected to be weighed differently according to which one is more relevant to perform the task. In other words, the encoding of numerosity should be stronger in the numerosity task compared to the duration task, and vice versa for duration. Overall, our results thus suggest that the ERP signals evoked by our stimuli reflect a perceptual aspect of magnitude encoding rather than decisional or mnemonic processes. In turn, the interaction between different magnitudes revealed by ERPs is thus likely to reflect a perceptual integration. This would be in line with a recent study from our group addressing the nature of magnitude integration (Togoli et al., 2022), providing evidence that this effect is perceptual in nature. However, as it is difficult to draw a clear distinction between perceptual and post-perceptual processes based on EEG results, our present considerations on this point remain speculative.

Finally, a limitation of the present study is that it only addressed the integration between two magnitude dimensions, numerosity and duration. According to the ATOM framework (Walsh, 2003), a true generalized magnitude processing stage should encompass several different magnitude dimensions, including space (i.e., for example as conveyed as the size of the stimuli), and potentially other dimensions that can be conceived as more vs. less, like the intensity of the stimuli (i.e., brightness). Future studies should thus address magnitude integration and its neural signature testing different pairs of dimensions, or even multiple dimensions at once, to reach more robust conclusions concerning the existence of a truly generalized processing stage mediating magnitude integration.

To conclude, our results show a specific latency window (∼360-460 ms) whereby visual evoked EEG responses are modulated by the interaction between numerosity and duration. The interaction effect within this window emerges irrespective of the task performed by participants, suggesting that it is not related to decisional or feature-based attentional processes. Additionally, activity within this window is successfully predicted by the magnitude integration bias measured behaviourally, suggesting a link between brain processing at this stage and the behavioural effect. We interpret brain activity within this latency window as a signature of a generalized magnitude processing stage mediating the integration of different dimensions. Overall, our results thus provide new evidence concerning the existence of a specific brain mechanism where information from different stimulus dimensions converges and is jointly processed.

## REFERENCES

Bendixen, A., Grimm, S., & Schröger, E. (2005). Human auditory event-related potentials predict duration judgments. Neuroscience Letters, 383(3), 284–288. https://doi.org/10.1016/j.neulet.2005.04.034

Bocincova, A., & Johnson, J. S. (2019). The time course of encoding and maintenance of task-relevant versus irrelevant object features in working memory. Cortex, 111, 196–209. https://doi.org/10.1016/j.cortex.2018.10.013

Borghesani, V., de Hevia, M. D., Viarouge, A., Pinheiro-Chagas, P., Eger, E., & Piazza, M. (2019). Processing number and length in the parietal cortex: Sharing resources, not a common code. Cortex, 114, 17–27. https://doi.org/10.1016/j.cortex.2018.07.017

Bueti, D., & Walsh, V. (2009). The parietal cortex and the representation of time, space, number and other magnitudes. Philosophical Transactions of the Royal Society of London. Series B, Biological Sciences, 364(1525), 1831–1840. https://doi.org/10.1098/rstb.2009.0028

Cai, Z. G., & Connell, L. (2015). Space–time interdependence: Evidence against asymmetric mapping between time and space. Cognition, 136, 268–281. https://doi.org/10.1016/j.cognition.2014.11.039

Cai, Z. G., & Wang, R. (2014). Numerical Magnitude Affects Temporal Memories but Not Time Encoding. PLoS ONE, 9(1), e83159. https://doi.org/10.1371/journal.pone.0083159

Cai, Z. G., Wang, R., Shen, M., & Speekenbrink, M. (2018). Cross-dimensional magnitude interactions arise from memory interference. Cognitive Psychology, 106, 21–42. https://doi.org/10.1016/j.cogpsych.2018.08.001

Casasanto, D., & Boroditsky, L. (2008). Time in the mind: Using space to think about time. Cognition, 106(2), 579–593. https://doi.org/10.1016/j.cognition.2007.03.004

Cheyette, S. J., & Piantadosi, S. T. (2020). A unified account of numerosity perception. Nature Human Behaviour, 4(12), 1265–1272. https://doi.org/10.1038/s41562-020-00946-0

Delorme, A., & Makeig, S. (2004). EEGLAB: an open source toolbox for analysis of single-trial EEG dynamics including independent component analysis. Journal of Neuroscience Methods, 134(1), 9–21. https://doi.org/10.1016/j.jneumeth.2003.10.009

DeWind, N. K., Adams, G. K., Platt, M. L., & Brannon, E. M. (2015). Modeling the approximate number system to quantify the contribution of visual stimulus features. Cognition, 142, 247–265. https://doi.org/10.1016/j.cognition.2015.05.016

Dormal, V., Seron, X., & Pesenti, M. (2006). Numerosity-duration interference: A Stroop experiment. Acta Psychologica, 121(2), 109–124. https://doi.org/10.1016/j.actpsy.2005.06.003

Ernst, B., Reichard, S. M., Riepl, R. F., Steinhauser, R., Zimmermann, S. F., & Steinhauser, M. (2017). The P3 and the subjective experience of time. Neuropsychologia, 103, 12–19. https://doi.org/10.1016/j.neuropsychologia.2017.06.033

Fornaciai, M., Brannon, E. M., Woldorff, M. G., & Park, J. (2017). Numerosity processing in early visual cortex. NeuroImage, 157. https://doi.org/10.1016/j.neuroimage.2017.05.069

Gao, Z., Yu, S., Zhu, C., Shui, R., Weng, X., Li, P., & Shen, M. (2016). Object-based Encoding in Visual Working Memory: Evidence from Memory-driven Attentional Capture. Scientific Reports, 6(1), 22822. https://doi.org/10.1038/srep22822

Harvey, B. M., Fracasso, A., Petridou, N., & Dumoulin, S. O. (2015). Topographic representations of object size and relationships with numerosity reveal generalized quantity processing in human parietal cortex. Proceedings of the National Academy of Sciences, 112(44), 13525–13530. https://doi.org/10.1073/pnas.1515414112

Harvey, B. M., Klein, B. P., Petridou, N., & Dumoulin, S. O. (2013). Topographic Representation of Numerosity in the Human Parietal Cortex. Science, 341(6150), 1123–1126. https://doi.org/10.1126/science.1239052

Javadi, A. H., & Aichelburg, C. (2012). When Time and Numerosity Interfere: The Longer the More, and the More the Longer. PLoS ONE, 7(7), e41496. https://doi.org/10.1371/journal.pone.0041496

Kleiner M, Brainard D, & Pelli D. (2007). What’s new in Psychtoolbox-3? Perception 36 ECVP Abstract Supplement.

Kononowicz, T. W., & Penney, T. B. (2016). The contingent negative variation (CNV): timing isn’t everything. Current Opinion in Behavioral Sciences, 8, 231–237. https://doi.org/10.1016/j.cobeha.2016.02.022

Lambrechts, A., Walsh, V., & van Wassenhove, V. (2013). Evidence Accumulation in the Magnitude System. PLoS ONE, 8(12), e82122. https://doi.org/10.1371/journal.pone.0082122

Lopez-Calderon, J., & Luck, S. J. (2014). ERPLAB: an open-source toolbox for the analysis of event-related potentials. Frontiers in Human Neuroscience, 8. https://doi.org/10.3389/fnhum.2014.00213

Macar, F., & Vidal, F. (2003). The CNV peak: An index of decision making and temporal memory. Psychophysiology, 40(6), 950–954. https://doi.org/10.1111/1469-8986.00113

Park, J., DeWind, N. K., Woldorff, M. G., & Brannon, E. M. (2016). Rapid and Direct Encoding of Numerosity in the Visual Stream. Cerebral Cortex, bhv017. https://doi.org/10.1093/cercor/bhv017

Pelli, D. G. (1997). The VideoToolbox software for visual psychophysics: Transforming numbers into movies. Spatial Vision, 10, 437–442.

Pouthas, V., Garnero, L., Ferrandez, A.-M., & Renault, B. (2000). ERPs and PET analysis of time perception: Spatial and temporal brain mapping during visual discrimination tasks. Human Brain Mapping, 10(2), 49–60. https://doi.org/10.1002/(SICI)1097-0193(200006)10:2<49::AID-HBM10>3.0.CO;2-8

Rammsayer, T. H., & Verner, M. (2014). The effect of nontemporal stimulus size on perceived duration as assessed by the method of reproduction. Journal of Vision, 14(5), 17–17. https://doi.org/10.1167/14.5.17

Skagerlund, K., Karlsson, T., & Träff, U. (2016). Magnitude Processing in the Brain: An fMRI Study of Time, Space, and Numerosity as a Shared Cortical System. Frontiers in Human Neuroscience, 10. https://doi.org/10.3389/fnhum.2016.00500

Temple, E., & Posner, M. I. (1998). Brain mechanisms of quantity are similar in 5-year-old children and adults. Proceedings of the National Academy of Sciences, 95(13), 7836–7841. https://doi.org/10.1073/pnas.95.13.7836

Togoli, I., Bueti, D., & Fornaciai, M. (under review). The nature of magnitude integration: contextual interference vs. active magnitude binding.

Togoli, I., Fornaciai, M., & Bueti, D. (2021). The specious interaction of time and numerosity perception. Proceedings of the Royal Society B: Biological Sciences, 288(1959), 20211577. https://doi.org/10.1098/rspb.2021.1577

Tonoyan, Y., Fornaciai, M., Parsons, B., & Bueti, D. (2020). Subjective time is predicted by local and early visual processing. BioRxiv. https://doi.org/10.1101/2020.10.30.362038

Walsh, V. (2003). A theory of magnitude: common cortical metrics of time, space and quantity. Trends in Cognitive Sciences, 7(11), 483–488. https://doi.org/10.1016/j.tics.2003.09.002

Watson, A. B. (1979). Probability summation over time. Vision Research, 19(5), 515–522. https://doi.org/10.1016/0042-6989(79)90136-6

Wichmann, F. A., & Hill, N. J. (2001). The psychometric function: I. Fitting, sampling, and goodness of fit. Perception & Psychophysics, 63(8), 1293–1313. https://doi.org/10.3758/BF03194544

Wiener, M., & Thompson, J. C. (2015). Repetition enhancement and memory effects for duration. NeuroImage, 113, 268–278. https://doi.org/10.1016/j.neuroimage.2015.03.054

Xu, F., & Spelke, E. S. (2000). Large number discrimination in 6-month-old infants. Cognition, 74(1), B1–B11. https://doi.org/10.1016/S0010-0277(99)00066-9

Xuan, B., Chen, X.-C., He, S., & Zhang, D.-R. (2009). Numerical magnitude modulates temporal comparison: An ERP study. Brain Research, 1269, 135–142. https://doi.org/10.1016/j.brainres.2009.03.016

Xuan, B., Zhang, D., He, S., & Chen, X. (2007). Larger stimuli are judged to last longer. Journal of Vision, 7(10), 2. https://doi.org/10.1167/7.10.2

Yates, M. J., Loetscher, T., & Nicholls, M. E. R. (2012). A generalized magnitude system for space, time, and quantity? A cautionary note. Journal of Vision, 12(7), 9–9. https://doi.org/10.1167/12.7.9

Zhang, M., Zhang, K., Zhou, X., Zhan, B., He, W., & Luo, W. (2021). Similar CNV Neurodynamic Patterns between Sub- and Supra-Second Time Perception. Brain Sciences, 11(10), 1362. https://doi.org/10.3390/brainsci11101362

